# Utilizing a cell culture based novel cellular thermal shift assay to understand the isoform-dependent thermal stability of ApoE variants

**DOI:** 10.64898/2026.06.09.730809

**Authors:** Rosemary J. Jackson, Steven Dierksmeier, Maryam Nishtar, Jonah Meltzer, Frieda Balduin, Bobby Beaumont, Zhanyun Fan, Eduard Sergienko, Steve Olson, Michael R. Jackson, Bradley T. Hyman

## Abstract

Apolipoprotein E (ApoE) is the primary genetic risk modifier of late-onset Alzheimer’s disease, with the ε4 allele increasing risk up to 15-fold relative to ε3. The structural differences between isoforms are thought to underlie their distinct effects on lipid transport, receptor binding, and disease risk. ApoE4 exhibits reduced thermodynamic stability compared to ApoE3, but prior characterisation has relied on purified recombinant protein, leaving open whether these differences are preserved in native cellular environments and how they relate to rare disease-associated variants. Here, we employed the cellular thermal shift assay (CETSA) and a bioluminescence-based thermal stability assay (BiTSA) to systematically characterise ApoE thermal stability across isoforms and variants. Using CETSA on brain tissue from humanised APOE knock-in mice and post-mortem human brain, we confirm that ApoE4 exhibits significantly reduced thermal stability compared to ApoE3 in native tissue, with this difference conserved across species despite variation in absolute melting temperatures. We developed BiTSA, which leverages a split-luciferase HiBiT tag to quantify soluble ApoE across a thermal gradient in living cells, providing a higher-throughput platform that faithfully recapitulates isoform stability differences. Applying BiTSA to rare AD-associated variants, we found that L28P exerts divergent, isoform-dependent effects, destabilising ApoE3 while paradoxically stabilising ApoE4—a finding supported by AlphaFold modelling revealing isoform-specific differences in helix 1 architecture. These results establish BiTSA as a robust cellular tool for ApoE variant characterisation and demonstrate that isoform background critically modulates the structural consequences of rare mutations.

## Introduction

Apolipoprotein E (ApoE) is a secreted glycoprotein that is the primary apolipoprotein in circulation in the central nervous system (CNS) and plays a critical role in lipid transport and cholesterol homeostasis^1^. In the brain, ApoE is primarily produced by astrocytes and microglia and is essential for the redistribution of lipids during neuronal development, maintenance, and repair^2^. The APOE gene is polymorphic, with three common alleles (ε2, ε3, and ε4) encoding three protein isoforms (ApoE2, ApoE3, and ApoE4) that differ at amino acid positions 112 and 158^3^. The ε4 allele is the strongest genetic risk factor for late-onset Alzheimer’s disease (AD), increasing risk 3-fold in heterozygotes and up to 15-fold in homozygotes compared to ε3 carriers ^4–6^. Conversely, the ε2 allele confers protection against AD^7,8^. ApoE isoform has been implicated in numerous disease-relevant processes, including neuroinflammation ^9,10^, amyloid-β and tau pathology^11,12^, synaptic dysfunction^13–15^, cerebrovascular impairment^16,17^, and mitochondrial and lysosomal stress ^18–20^.

ApoE is a 299-amino acid protein consisting of two independently folded domains: an N-terminal domain (residues 1-191) containing a four-helix bundle that harbors the LDL receptor binding region, and a C-terminal domain (residues 210-299) responsible for lipid binding^20–23^. These domains are connected by a flexible hinge region. The N-terminal four-helix bundle is critical for receptor interactions, while the C-terminal domain mediates lipid association and protein oligomerization^23^. Recent single-molecule FRET and molecular dynamics studies have demonstrated that ApoE4 exhibits extensive conformational heterogeneity in both lipid-free and lipid-bound forms, being far more disordered and extended than previously thought^24^. This structural plasticity, with the protein adopting closed, open, and extended conformations, may underlie the functional differences between isoforms. Recent cryo-EM studies have provided important structural insights, demonstrating that ApoE secreted by astrocytes forms antiparallel dimers in discoidal lipoproteins, adopting a “double-belt” configuration that has important implications for understanding its function in health and disease ^25^.

The three ApoE isoforms differ at only two amino acid positions, with ApoE2 having a cysteine at both positions 112 and 158, ApoE3 having a cysteine at 112 and arginine at 158, and ApoE4 having arginine at both positions^3^. These seemingly minor differences have profound effects on protein structure and function. The arginine at position 112 in ApoE4 is predicted to promote an interaction between the N- and C-terminal domains (domain interaction) that is absent in ApoE3 and ApoE2 ^23,26,27^. This domain interaction alters the protein’s lipid binding properties, receptor binding affinity, and stability^24,28–30^. Lipidation of ApoE, which occurs in vivo, protects the protein from aggregation, potentially impacting its physiological function^31^.

Structural studies have revealed that ApoE4 exhibits reduced thermodynamic stability compared to ApoE3, with the four-helix bundle of ApoE4 being more susceptible to denaturation^28,32,33^. This reduced stability has been attributed to the domain interaction unique to ApoE4, which places strain on the N-terminal helix bundle^26^. Hydrogen-deuterium exchange mass spectrometry (HDX-MS) studies have shown that the N-terminal helix of ApoE4 begins at a later point in the sequence compared to ApoE3, resulting in a shorter and potentially less stable helical region^33^. Cross-linking mass spectrometry combined with molecular modeling has further characterized the lipidated ApoE4 structure and its receptor binding mechanism, revealing how the protein adapts its conformation for recognition by the LDL receptor^34^.s

Recently, several mutations have been identified that either alter, or are predicted to alter, the thermal stability of ApoE. These include both disease-associated variants and protective variants. The APOE3-Christchurch variant (R136S) was identified in a patient who remained cognitively intact despite carrying a presenilin-1 mutation that typically causes early-onset AD^35^. Similarly, the APOE3-Jacksonville (V236E) variant and APOE4-R251G mutations have been associated with reduced risk of dementia and AD^36,37^ while APOE3-R145C has been shown to increase the risk of AD^38^. Understanding how these variants affect protein stability may provide insights into the mechanisms underlying their protective or pathogenic effects.

Protein thermal stability is a fundamental biophysical property that reflects the overall structural integrity and folding state of a protein. Thermal shift assays are widely used to measure protein stability by monitoring the temperature-dependent unfolding of proteins^39^. In conventional protein thermal shift (PTS) assays, fluorescent dyes such as SYPRO Orange bind to hydrophobic residues that become exposed as proteins unfold, generating a fluorescence signal that can be used to determine the melting temperature (Tm). However, ApoE presents unique challenges for conventional PTS approaches. Lipid-free ApoE readily self-associates through hydrophobic interactions, forming oligomers and aggregates that complicate thermal stability measurements ^40,41^. Furthermore, ApoE contains numerous hydrophobic residues on its surface that are involved in lipid binding, which can interact with hydrophobic dyes even in the native state, confounding the interpretation of thermal shift data. The Cellular Thermal Shift Assay (CETSA) offers an alternative approach that circumvents many of these limitations by measuring protein stability in the native cellular environment^42,43^. In CETSA, cells or tissue lysates are heated to different temperatures, and the amount of soluble (non-aggregated) protein remaining is quantified by immunoblotting or other detection methods. This approach allows proteins to be assessed in their physiological context, including relevant post-translational modifications, binding partners, and lipid associations ^44^.

In this study, we employed CETSA and a novel high-throughput bioluminescence-based thermal stability assay (BiTSA)^45^ to systematically characterize the thermal stability of ApoE isoforms and variants. We demonstrate that ApoE4 exhibits reduced thermal stability compared to ApoE3 in both mouse and human brain tissue, and that this difference is recapitulated in our cell-based BiTSA system. We further show that rare ApoE variants associated with altered AD risk have distinct effects on thermal stability, providing new insights into the structure-function relationships of this important protein.

## Methods

### Animals

*APOE* knock-in mice expressing human *APOE*ε2, *APOE*ε3 or *APOE*ε4 in the locus of the murine *Apoe* gene^46^ were used in this experiment as were C57BL/6NTac. 10–12-month-old mice were euthanized by CO_2_ asphyxiation, transcardially perfused with ice-cold PBS and their brain tissue was collected and snap frozen. Mice were group housed in an animal facility with controlled conditions of temperature and humidity, 12/12-h dark/light cycle and ad libitum access to food and water. All animal experiments were approved by the Massachusetts General Hospital Subcommittee on Research Animal Care following the guidelines set forth by the National Institutes of Health Guide for the Care and Use of Laboratory Animals (no. 2006n000178). Both sexes were used.

African green monkey tissue was obtained from the Davidson lab and was naïve of manipulation. Both sexes were used. All animal procedures were approved by the CHOP Institutional Animal Care and Use Committee.

### Human brain collection

Human brain tissue was obtained from the Massachusetts Alzheimer Disease Research Center Tissue Bank. Tissue was dissected at the time of autopsy and immediately snap frozen with powdered dry ice. Postmortem intervals were less than 24 h. All sample collection was obtained under a research protocol approved by the Massachusetts General Hospital IRB.

### CETSA

Brain tissue was homogenized in 10 volumes by weight of ice-cold TBS with protease and phosphatase inhibitors using a Dounce homogenizer. The homogenate was then spun at 10 000*g* for 10 min and the supernatant (TBS-soluble fraction) was collected for western blot. Protein concentration was determined using a BCA assay. Lysates were diluted 0.75mg/ml where required and 50ul of sample were placed into each of eight 0.2ml PCR tubes. Samples were then heated in an Eppendorf 96 well X50H (Eppendorf) in a thermal gradient from 37°C to 62°C degrees for 15min followed by a rapid cooling to 4°C for 5 minutes.

#### Western blot

Samples were then removed from the heat block and centrifuged at 20,000G for 20 minutes. 13ul of Supernatant was combined with 2ul of NuPAGE Sample Reducing Agent (10X) (Invitrogen, #B0009) and 5ul of NuPAGE LDS Sample Buffer (4X) (Invitrogen, #NP0007) and heated to 95°C for 5 minutes before 13ul of this was loaded onto a NuPage 4-12% Bis-Tris gels and in MES buffer.

Proteins were electrotransferred onto nitrocellulose membrane at 30 V for 1.5 h using the XCell II™ Blot Module system in tris-glycine transfer buffer. Membranes were incubated in blocking buffer (Li-Cor Biosciences) diluted 1:1 TBS for 1 h to reduce background staining. Membranes were then incubated with primary antibody; rb anti-ApoE (Novus biologicals, NBP1-31123), Mouse Anti-HiBiT Antibody (30E5) (Promega, #CS2006A01) diluted in blocking buffer with added 0.1% Tween-20 overnight at room temperature while shaking. Membranes were then washed and incubated with the appropriate 680 IR dye secondary antibodies (Li-Cor Biosciences). The membranes were imaged using Odyssey infrared imaging system, and analyzed using Odyssey software.

### Plasmid prep

Mammalian expression plasmids based on pcDNA3.1 containing mouse ApoE, monkey ApoE, ApoE3, or ApoE4 with the HiBiT tag on the N-terminus were custom prepared by Biomatik (Kitchener, ON, Canada). Maxipreps were generated using the Qiagen maxiprep kit (Qiagen) according to manufacturer instructions.

Variants were introduced into the backbone of ApoE indicated using the QuikChange II kit (agilent) and according to manufacturers instructions. Minipreps were generated using the Qiagen miniprep kit (Qiagen) according to manufacturer instructions.

The sequence of all plasmids was confirmed by rapid plasmid sequencing at the Massachusetts General Hospital DNA Core.

### Cell culture

HEK-293 cells were maintained at 37°C in 5% CO_2_ in Dulbecco’s modified Eagle medium (DMEM), 10% fetal bovine serum (FBS) 1% Glutamax and 1% penicillin–streptomycin. Cells were plated into 12 well plates at 50k cells per well and allowed to recover overnight. The following days cells were transfected using 1.2ug of DNA and 4ul of Lipofectamine 2000 (Invitrogen) per well. 4-6 hours following transfection cells media was removed and exchanged for OMEM with 1% penicillin–streptomycin and cultured at 37°C in 5% CO_2_ for 16-20 hours

### BITSA

Media was removed and cells were incubated in 100ul of enzyme free Cell Dissociation Buffer (Gibco) for 5 minutes after which 650ul of OMEM was added. Cells were pipetted several times to generate a single cell suspension and then 10ul of cell suspension was added to each 64 wells of an opaque conical 384 well plate (bio-rad). Samples were then heated in an Eppendorf 384 well X50H (Eppendorf) in a thermal gradient from 36°C to 61°C degrees for 8min followed by a rapid cooling to 4°C for 3 minutes. The thermal gradient runs top to bottom so 16 points of the thermal curve were assessed in technical quadruplet. 10ul of detection mix from the Nano-Glo HiBiT Lytic Detection System (Promega) was added and the plate was incubated at room temp in the dark for 15 minutes. Luminescence signals were read in by Victor 3V luminescence plate reader (Perkin-Elmer).

### Generating astrocytes for BITSA

ApoE-knockout mouse astrocytes^47^ were transduced with lentivirus according to the Lentivirus Transduction Protocol by Vectorbuilder. HiBiT-tagged ε2, ε3, and ε4 lentivirus was added to the cells, adapted to a Multiplicity of Infection (MOI) of 5 along with 6ug polybrene per well. DMEM was used for a media change after 20h and subsequent cell maintenance. Successfully transduced astrocytes were selected in a 2-week incubation period with 8ul Hygromycin B (Invitrogen, #10687010) per well. (ScienCell #1801). Successful transduction was measured following instructions of the Nano-Glo HiBiT Lytic Detection System measured in the Perkin-Elmer Victor 3V luminescence plate reader.

### Alpha-fold

Three-dimensional protein structures of apolipoprotein E isoforms (ApoE2, ApoE3, and ApoE4) and their variants were predicted using the AlphaFold Server powered by AlphaFold 3 and developed by Google DeepMind^48^. The canonical full-length amino acid sequence of human APOE (P02649) was used as the reference sequence for ApoE3. Isoform-specific sequences were derived by introducing the defining single amino acid substitutions: ApoE2 (Cys112, Cys158), ApoE3 (Cys112, Arg158), and ApoE4 (Arg112, Arg158), relative to the canonical sequence. Variant sequences (e.g., E96K, R114C) were generated by manual substitution at the relevant positions. Each sequence was submitted individually to the AlphaFold Server as a single protein chain input with default parameters. Predicted structures were downloaded in PDB format and visualised using PyMol (Schrödinger). Predicted Local Distance Difference Test (pLDDT) scores were used to assess per-residue confidence, with scores above 70 considered reliable for structural interpretation, as recommended.

### Statistics

For CETSA and BiTSA data was normalized across runs by defining the lowest temperature as 100% signal and the highest temperature as 0% signal. A sigmoidal Boltzmann curve, with Bottom equal to 0, Top equal to 100, V50 > 0, and slope < 0 was then fit to the data and the V50 and slope were recorded. Analysis of non-normalized data yielded results that did not differ statistically from those obtained with normalized data.

Normality was assessed using the Shapiro-Wilk test. One-way ANOVA’s were used to assess the V50 and slope differences with Dunnett’s multiple comparisons test to the ApoE backbone or ApoE4 as indicated or using Tukey’s multiple comparisons test. Human brain sample V50s were assessed using a students t-test.

## Results

### Thermal stability of ApoE isoforms in brain tissue

To assess the intrinsic thermal stability of apolipoprotein E (ApoE) isoforms in their native biochemical environment, we performed cellular thermal shift assays (CETSA)^42^ on brain tissue from humanized APOE knock-in mice expressing either human APOE2, APOE3 or APOE4^46^. Brain homogenates were subjected to a temperature gradient, and the amount of soluble ApoE remaining at each temperature was quantified by immunoblotting. We observed that ApoE4 exhibited a significantly lower melting temperature (Tm) compared to ApoE3, indicating reduced thermal stability of the ApoE4 isoform in the context of brain tissue (Figure 1A-B) [F (2, 12) = 13.70 (post-hoc tukeys ApoE3 to ApoE4 p=0.0159)].

**Figure 1.**
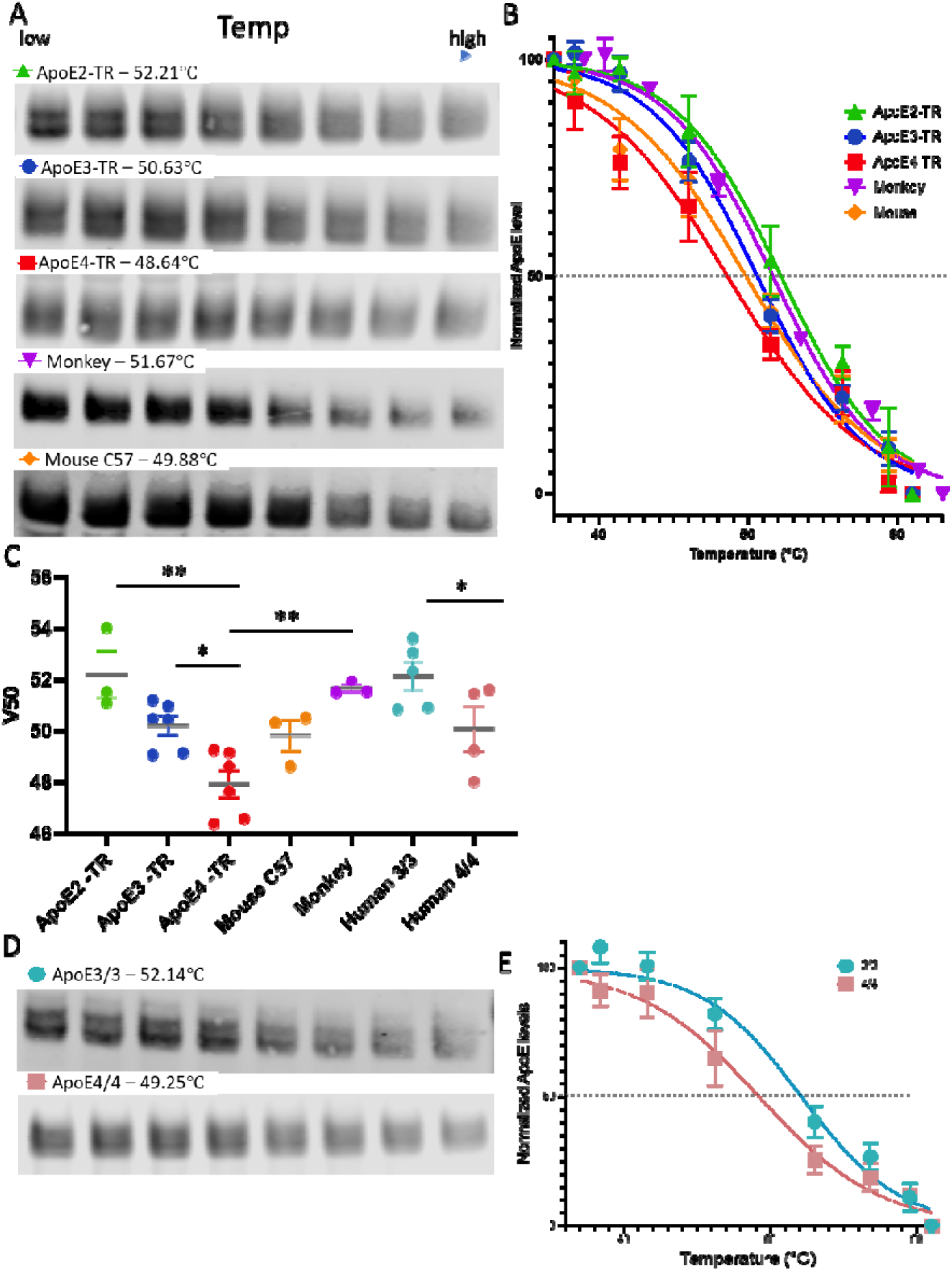
Thermal stability of ApoE isoforms in brain tissue. (A) Representative western blots of ApoE from TBS-soluble brain lysate from different models subjected to a temperature gradient (37-70°C). Mice with different humanized ApoE isoforms show different melting points (V50, the temperature at which 50% of protein remains soluble, indicated next to model name). (B) Modified Boltzmann sigmoidal curve of melting temperatures from model system CETSA. (C) V50 values for different APOE genotypes from CETSA analysis. (D) CETSA analysis of APOE in human post-mortem brain tissue from APOE3/3 and APOE4/4 individuals. (E) Modified Boltzmann sigmoidal curve of melting temperatures from human brain CETSA.

Comparison across species demonstrated that endogenous mouse ApoE exhibited a melting temperature comparable to human ApoE3, while monkey ApoE melted at a temperature similar to that of ApoE2 [F (6, 23) = 7.453 (post-hoc tukeys ApoE4 to Monkey p=0.0045)]. Given that both monkey and mouse ApoE have an arginine at position 112, this suggests that the thermal stability of the protein is linked to whole protein domain interactions rather than individual amino acid differences alone, as we expected. The reduced stability of ApoE4 relative to ApoE3 is consistent with previous biophysical studies using purified recombinant protein^28,32^.

To determine whether these isoform-dependent stability differences extend to the human brain, we performed CETSA on post-mortem human brain tissue from APOE3/3 and APOE4/4 homozygous individuals. Consistent with our findings in humanized mice, human ApoE4 displayed reduced thermal stability compared to ApoE3 (Figure 1C-D)[ t=2.071, df=7, p=0.0386]. Notably, while the delta between ApoE3 and ApoE4 is the same in both contexts, the absolute values are different, possibly reflecting differences in ApoE binding partners or lipid associations between mouse and human brain. These results establish that the thermal stability differences between ApoE isoforms observed with purified proteins are maintained in the complex environment of brain tissue, where ApoE exists in association with lipids and other binding partners.

### Development of a high-throughput in vitro assay to measure ApoE thermal stability

While CETSA provides a powerful means to assess protein stability in native tissues, it is inherently low-throughput and requires substantial amounts of biological material. To enable systematic characterization of ApoE variants, we developed a bioluminescence-based thermal stability assay (BiTSA) that allows rapid, quantitative measurement of protein thermal stability in living cells.

In this assay, ApoE variants were expressed as fusion proteins containing an N-terminal HiBiT tag ^45^ (Supplementary Figure 1A-B). Upon cell lysis in the presence of LgBiT, the complemented NanoLuc luciferase produces a luminescent signal proportional to the amount of soluble, properly folded ApoE. By measuring luminescence across a temperature gradient, we can determine the apparent melting temperature of each ApoE variant with high precision and throughput (Figure 2A).

**Figure 2.**
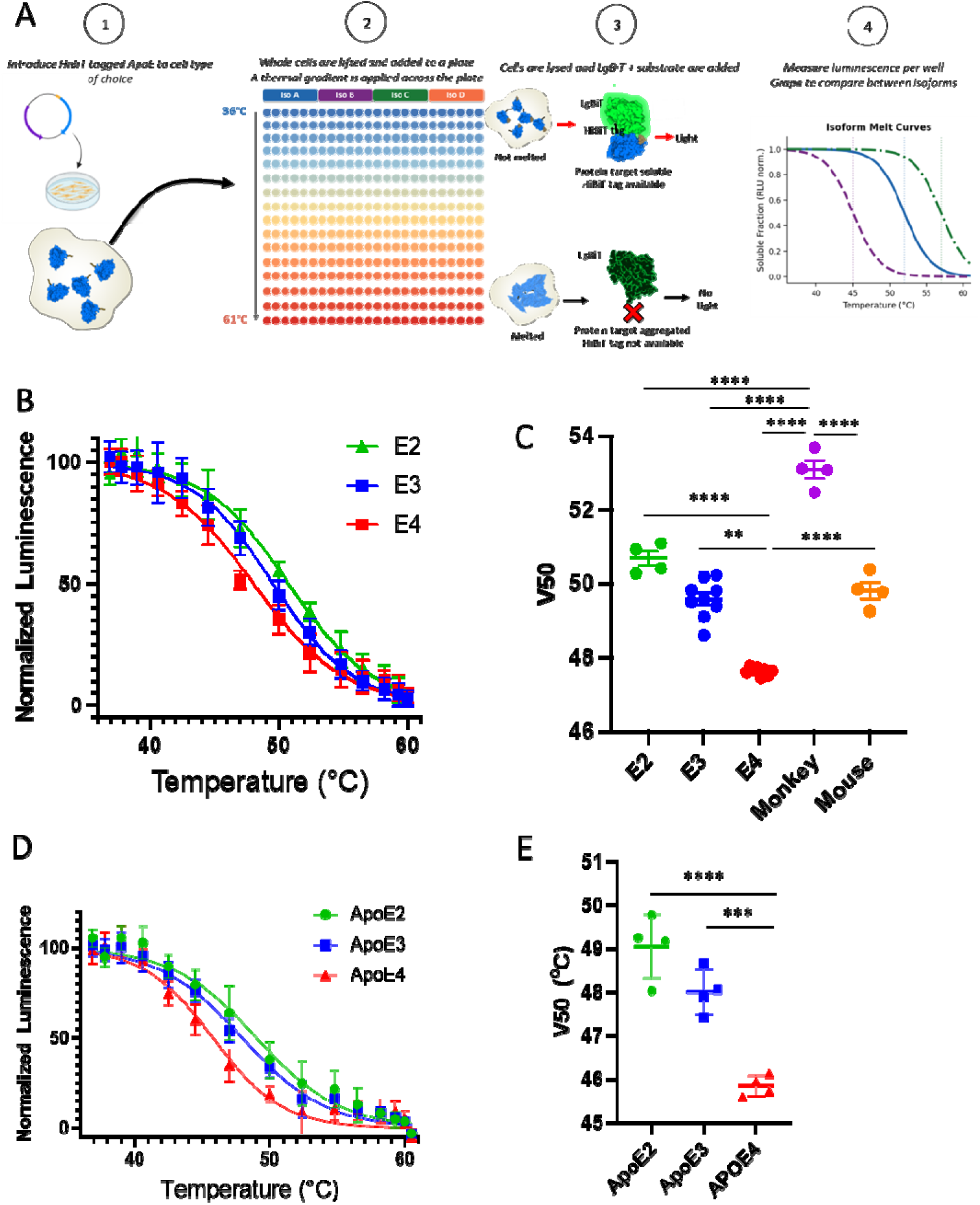
Development and validation of BiTSA for ApoE thermal stability measurement. (A) Schematic of the BiTSA system. (B) BiTSA analysis of ApoE isoforms in HEK293 cells, (C) Quantification of melting temperatures in HEK293 cells. (D) BiTSA analysis of ApoE isoforms in mouse immortalized astrocytes, (E) Quantification of melting temperatures in mouse immortalized astrocytes (*p < 0.05, **p < 0.01, ****p < 0.0001, tukeys post-hoc)

Using this system in HEK293 cells, we observed that the relative thermal stabilities of ApoE isoforms mirrored those observed in brain tissue CETSA experiments. ApoE4 consistently exhibited a lower Tm than ApoE3 (Figure 2B-C)[ F (4, 24) = 125.3, (Tukey’s post-hoc ApoE3 to ApoE4 p<0.0001)]. The delta of the stability difference between ApoE3 and ApoE4 (∼2-3°C) was consistent with previous reports using purified recombinant protein^28,32^, although again the absolute values differed slightly, likely reflecting differences in system complexity and the presence of cellular binding partners.

To further validate the robustness of this assay in a more physiologically relevant cell type, we expressed ApoE variants in immortalized human astrocytes. The stability differences between isoforms were maintained in this glial cell context, supporting the use of BiTSA as a reliable method for assessing ApoE thermal stability (Figure 2D-E).

### Testing the thermal stability of rare disease-associated ApoE variants

We next used this system to examine whether ApoE thermal stability correlates with AD risk profile. We tested ApoE3-Christchurch^35^, ApoE3-R145C^38^, ApoE3-Jacksonville^36,37^, and ApoE4-R251G^37^. ApoE3-Christchurch and ApoE3-Jacksonville have both been shown to be protective when compared with ApoE3^35–37^ In contrast, ApoE3-R145C has been associated with increased risk when compared with ApoE3, and ApoE4-R251G has been shown to decrease the risk of ApoE4 to be similar to that of ApoE3^37,38^ (Figure 3A-B).

**Figure 3.**
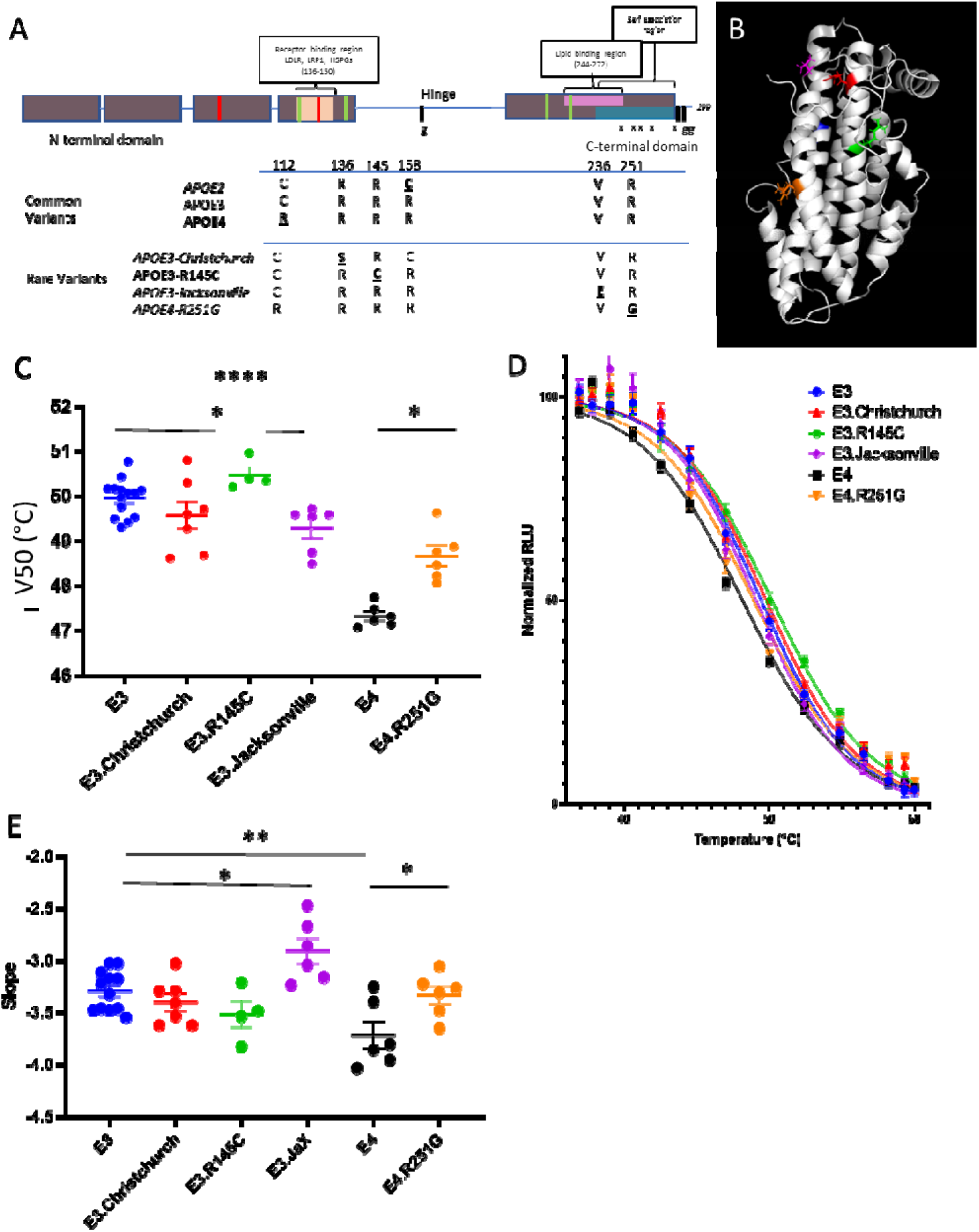
Thermal stability of disease-associated ApoE variants. (A) Schematic of rare ApoE variants indicating if the variant is associated with reduced AD risk (green, italics) or increased AD risk (red, bold) and the location of the mutation within the protein. (B) Annotated side chains of ApoE from PDB 2L7B^40^. (C) V50 values for different ApoE variants from BiTSA analysis. (D) Modified Boltzmann sigmoidal curve of melting temperatures from BiTSA. (E) Slope of the curve values for different ApoE variants from BiTSA. (*p < 0.05, **p < 0.01, ****p < 0.0001, Dunnet’s post-hoc to ApoE backbone)

We introduced these rare variants into the ApoE backbone with which they are associated in the population and found that while ApoE3-Christchurch and ApoE3-R145C had no impact on the thermal stability, ApoE3-Jacksonville caused a slight destabilization of the protein (Figure 3C-D)[ F(5, 37) = 28.98 (Dunnett’s ApoE3 vs ApoE3_Jacksonville p=0.0446)]. This corresponded to a similarly significant reduction in the width of the slope, which could be due to the reduced propensity of ApoE3-Jacksonville to dimerize or oligomerize^36^ (Figure 3E)[ F (5, 35) = 7.419 (Dunnets ApoE3 vs ApoE3_Jacksonville p=0.0142, ApoE3 vs ApoE4 p=0.0056). ApoE4-R251G, however, caused a slight but significant increase in the thermal stability when compared with ApoE4 (Dunnets ApoE4 vs ApoE4_R251G p=0.0003) (Figure 3C-D). Collectively, these results indicate that thermal stability does not directly correlate with disease risk status, suggesting that mutations in the N-terminal receptor binding region likely act via mechanisms other than altered protein stability, such as heparan sulfate binding^35^.

### Thermal stability analysis of additional rare ApoE variants

To further explore the relationship between ApoE structure and thermal stability, we used this technique to examine a panel of rare ApoE variants (Figure 4A) that have been identified in human populations and associated with various phenotypes including altered lipid metabolism and dementia risk.

**Figure 4.**
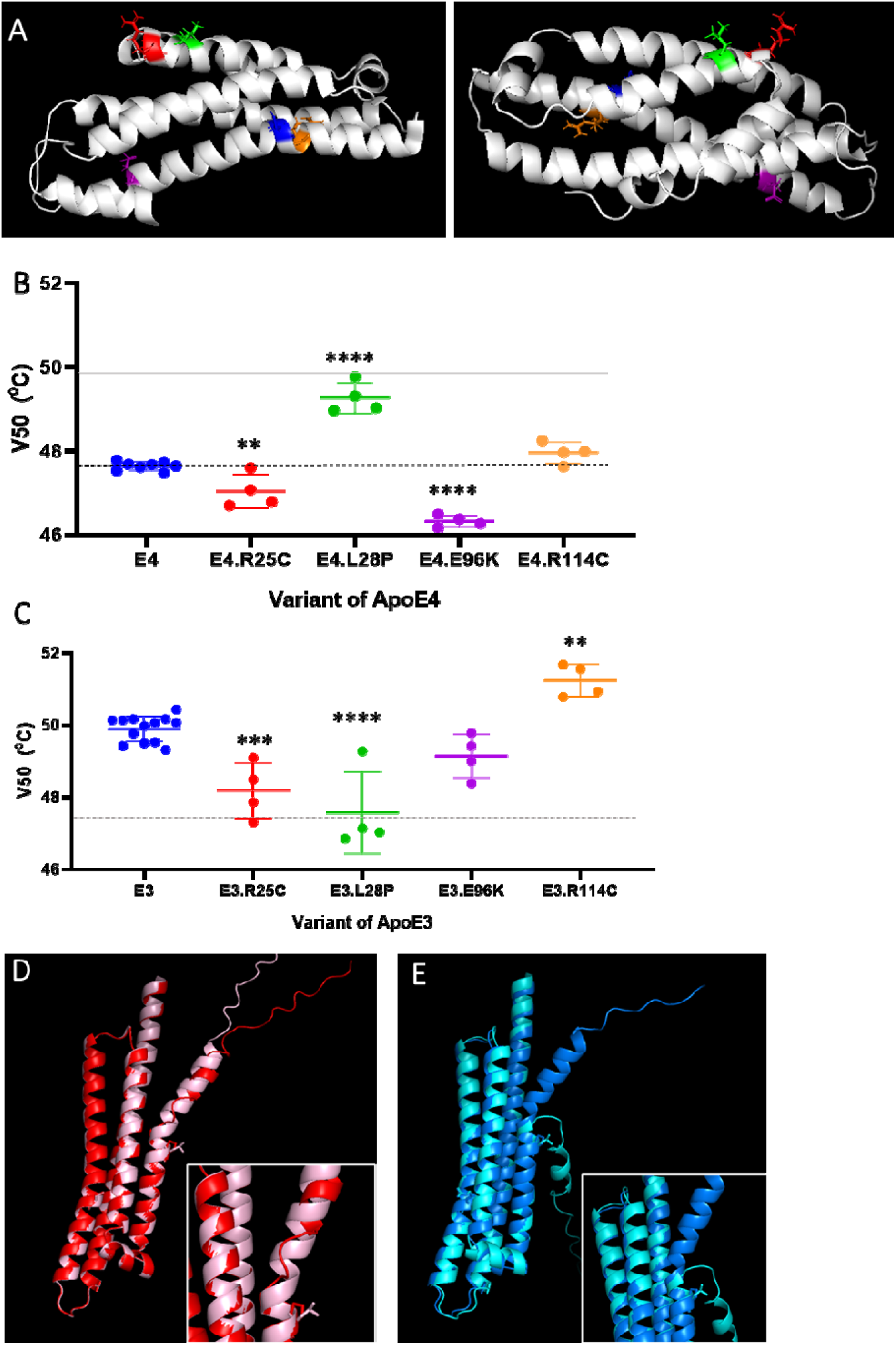
Thermal stability analysis of ApoE variants. (A) Annotated side chains of the N-terminal domain of ApoE from PDB 2L7B in two orientations. (B) BiTSA analysis of the rare variants in the ApoE3 backbone. (C) BiTSA analysis of the rare variants in the ApoE4 backbone. (D) AlphaFold model of ApoE3-L28P showing disruption of helix 1 structure; pink represents native ApoE3 and dark red represents ApoE3-L28P. (E) AlphaFold model of ApoE4-L28P showing restoration of helical structure; cyan represents native ApoE4 and blue represents ApoE4-L28P. (*p < 0.05, **p < 0.01, ***p < 0.001, ****p < 0.0001, Dunnetts pos-hoc to ApoE backbone)

R25C (ApoE-Kyoto)^49^ and R114C^50^ are both arginine-to-cysteine substitutions located in the N-terminal helix bundle domain. R25C was first identified in Japanese patients with type III hyperlipoproteinemia and has been associated with altered lipid binding properties^49^. R114C is located near the critical position 112 that distinguishes ApoE4 from ApoE3, and substitutions at this position might be expected to influence the domain interaction characteristic of ApoE4.

In our BiTSA analysis, R25C caused modest destabilization when introduced into both the ApoE3 and the ApoE4 backbone (Figure 4B-C), while R114C showed a modest but significant stabilizing effect in ApoE3 but not ApoE4. [APOE4 ANOVA F (4, 19) = 75.88, (Dunnets ApoE4 vs ApoE4_R25C p=0.0075), APOE 3 ANOVA F (4, 24) = 24.50 (Dunnets ApoE3 vs ApoE3_R25C p=0.0002, ApoE3 vs ApoE3_R114C p=0.0029)]. These divegent responses suggest that amino acid substitutions have differential effects depending on the local environment of the protein and the potential domain-domain interactions interrupted. The parallel effect of R25C^51^ across isoforms suggests that this region is structurally similar between ApoE3 and ApoE4, whereas the divergent effects of R114C highlight differences in the region surrounding position 112.

The E96K variant has been associated with increased risk of dementia and AD in a large Danish cohort study^52^. This glutamate-to-lysine substitution introduces a positive charge in the N-terminal domain, potentially disrupting local electrostatic interactions. Intriguingly, E96K showed no impact on the thermal stability in ApoE3 although it showed significant destabilization of ApoE4 [Dunnets ApoE4 vs ApoE4_E96K p<0.0001], which is notable given that the change from an acidic glutamic acid to a basic lysine might be expected to disrupt protein structure for both isoform backbones (Figure 4B-C).

The L28P mutation, also known as ApoE4-Pittsburgh or ApoE4-Freiburg, was initially identified in populations with altered lipid profiles and has a mixed association with AD with some studies indicating an increased risk and others no risk.^53–56^. Leucine 28 is located in the first helix of the N-terminal four-helix bundle, and its substitution with proline would be expected to disrupt the helical structure due to proline’s unique conformational constraints.

Strikingly, we observed a divergent, isoform-dependent effect of L28P on thermal stability. Introduction of L28P into ApoE3 resulted in significant destabilization, consistent with the expected disruption of helical structure (Figure 4C) [Dunnets ApoE3 vs ApoE3_L28P p<0.0001]. However, when L28P was introduced into ApoE4, we observed a paradoxical stabilization of the protein (Figure 4B)[Dunnets ApoE4 vs ApoE4_L28P p<0.0001]. This unexpected finding suggests that the structural context provided by the ApoE4 backbone, particularly the domain interaction, may compensate for or even benefit from the conformational changes induced by the proline substitution.

To gain structural insight into this divergent response, we generated AlphaFold^48^ models of APOE3 and APOE4 carrying the L28P mutation. In the ApoE3-L28P model (Figure 4D), the proline substitution introduces a pronounced kink in helix 1 in four of five predicted models, disrupting the regular helical geometry and potentially destabilizing inter-helical packing within the four-helix bundle. In contrast, the ApoE4-L28P model (Figure 4E) shows that the helical structure in this region is largely restored (again in four of five models), possibly because the altered domain interaction in ApoE4 provides compensatory stabilizing contacts.

This observation aligns with HDX-MS studies demonstrating that the N-terminal helix of ApoE4 begins at a later point in the sequence compared to ApoE3, resulting in inherent differences in the helical structure of this region between isoforms^33^. The L28P mutation may therefore have different structural consequences depending on the pre-existing helical architecture of each isoform.

## Discussion

In this study, we demonstrate that both CETSA^42^ and the BiTSA^45^ assay are effective methods for assessing the thermal stability of ApoE isoforms and variants. Our findings confirm that ApoE4 exhibits reduced thermal stability compared to ApoE3 in both mouse and human brain tissue, consistent with previous biophysical studies using purified recombinant protein^28,32,33^. Importantly, we show that these stability differences are maintained in the complex cellular environment where ApoE exists in association with lipids and other binding partners. Notably, while the delta between ApoE3 and ApoE4 was conserved across contexts, there were differences in absolute values between contexts, which almost certainly indicates the presence of different ApoE binding partners or lipids. Further studies will be required to identify these binding partners and to determine their structural and functional consequences. It is intriguing that in the context of both mouse brain and mouse immortalized astrocytes the Tms were ∼1°C lower than the corresponding human context.

Our findings reveal that mutations in certain regions of ApoE are more likely to impact thermal stability than others. It is important to note that a change in thermal stability can be interpreted as evidence of altered conformational and structural stability; thus these findings provide a direct window into how specific mutations perturb ApoE structure in a physiologically relevant context. Variants affecting helix 1, such as R25C and L28P, and the C-terminal domain, R251G and Jacksonville, altered thermal stability, whereas variants in other regions showed more variable effects. This is consistent with the importance of the helical bundle and domain-domain interactions for overall protein stability and suggests that thermal stability may be a particularly sensitive indicator of structural perturbation in these regions ^22,24,33,41,51^. Of interest also are the variants that did not impact stability in either variant namely ApoE3_R145C and ApoE3_Christchurch which indicate that these mutations are likely causing their effect due to other properties such as receptor or heparin binding^35^.

Interestingly, some mutations had consistent effects on ApoE stability regardless of isoform backbone (e.g., R25C), while others showed striking isoform-dependent effects (e.g., L28P, R114C, E96K). The parallel destabilization caused by R25C suggests that this region of the protein is structurally similar between ApoE3 and ApoE4. In contrast the isoform-specific response suggests that the local electrostatic environment around residues 96 and 114 differ meaningfully between the two isoforms, likely as a consequence of the ApoE4-specific domain interaction driven by Arg112^57^. R114C caused a modest but significant stabilization of ApoE3, while E96K caused a de-stabilization of ApoE4. It is possible that in ApoE3, where Cys112 is already present, the introduction of a second cysteine at position 114 may alter local side-chain packing, permit a favorable intra-helical interaction, or may increase the formation of cystine dimers. Where-as this domain interaction allows the charge reversal introduced by E96K, from the negatively charged glutamate to the positively charged lysine, to be tolerated in ApoE3, where the surrounding helix is more flexible or electrostatically buffered, but is destabilizing in ApoE4, where the Arg112-mediated domain interaction renders the local environment less able to accommodate the additional positive charge without disrupting inter-helical electrostatic contacts and reducing overall bundle stability. More studies are needed to understand the impact of modifying helix 3 side chains in ApoE3 and ApoE4.

The paradoxical stabilization of ApoE4 by L28P merits particular attention. Our findings appear to contrast with previous work by^58,59^,who reported that L28P destabilizes ApoE4 in studies using purified recombinant protein. Several factors may account for this discrepancy. First, our BiTSA assay measures thermal stability in the cellular context, where ApoE is associated with lipids and other binding partners that may modulate the effects of mutations, recent studies have shown that lipidation of ApoE protects it from aggregation^31^. Second, post-translational modifications present in cells but absent in recombinant protein may influence the structural consequences of the L28P mutation^60,61^. Third, the oligomeric state of ApoE differs between cellular and in vitro conditions; recent single-molecule studies have shown that monomeric ApoE4 adopts conformations distinct from those observed in oligomeric forms^24^. These considerations underscore the importance of studying ApoE variants in physiologically relevant contexts.

It is important to acknowledge the limitations of thermal stability assays, including both CETSA and BiTSA. These assays measure the temperature at which proteins aggregate or become insoluble, which is influenced by multiple factors beyond intrinsic thermodynamic stability, including protein concentration, aggregation propensity, and interactions with other cellular and media components. ApoE presents particular challenges due to its well-documented tendency to self-associate and form oligomers^40,62^. Lipid-free ApoE readily self-associates through hydrophobic interactions, and this aggregation-prone behavior can complicate the interpretation of thermal stability data^41,63^. Despite these limitations, the consistency of our findings across different experimental systems (CETSA in brain tissue and BiTSA in cultured cells) supports the validity of our conclusions. It is also important to note that analysis of additional melt curve parameters, such as slope, may provide further insight into the oligomeric and aggregation behaviour of ApoE variants. Crucially, while a detected difference in thermal stability between variants can be interpreted as positive evidence of a change in conformational and structural stability, the converse, however, is not necessarily true: the absence of a thermal stability difference cannot be taken as evidence that the protein is conformationally unperturbed, as structural changes that do not alter aggregation temperature would not be detected.

A key strength of this study is that our analysis began with ApoE from human brain tissue, establishing the physiological relevance of our findings before moving to higher-throughput cellular systems. The concordance between brain tissue CETSA and cellular BiTSA results validates the use of BiTSA for systematic characterization of ApoE variants. This approach enabled us to examine a larger panel of variants than would be feasible with tissue-based methods alone.

We also need to understand what the rare variants that did not impact melting temperature do at the molecular level. Thermal stability is just one aspect of protein function, and variants that do not affect stability may still have important effects on lipid binding, receptor interactions, or other functional properties. Future studies combining thermal stability measurements with functional assays will be needed to fully characterize the consequences of ApoE variants.

In conclusion, this study establishes BiTSA as a robust and scalable approach for assessing the thermal stability of ApoE variants in a cellular context. Our findings reveal complex relationships between ApoE sequence, isoform background, and thermal stability that have implications for understanding the structural basis of ApoE-associated disease risk. The isoform-dependent effects of certain mutations highlight the importance of considering genetic background when evaluating the consequences of rare variants, both for mechanistic understanding and for clinical interpretation of genetic findings.

## Author contributions

R.J.J. was involved in conceptualization, data curation, formal analysis, funding acquisition, investigation, methodology, project administration, resources, software, supervision, validation, visualization, writing – original draft, and writing – review & editing. S.D., M.N., and F.B. were involved in investigation and methodology. J.M. was involved in investigation, methodology, and supervision. B.B. was involved in investigation and validation. Z.F. was involved in investigation and resources. E.S. was involved in formal analysis, funding acquisition, resources, and software. S.O. was involved in funding acquisition. M.R.J. and B.T.H. were involved in funding acquisition, supervision, and writing – review & editing.

## Acknowledgements and funding

The authors acknowledge funding from Alzheimer’s Drug Discovery Foundation (ADDF) to R. Jackson, from Alzheimer’s San Diego to M. Jackson and grants from the National Institutes of Health (NIH) T32AG000222-27 (R Jackson and B. Hyman), 5R56AG080525 (M. Jackson and B. Hyman), and 1R01AG086914 (M. Jackson). We would like to acknowledge funding from the JPB Foundation, and to the Mass ADRC (P30AG062421). R Jackson would also like to acknowledge the help, support and guidance from the Harrington foundation.

## Dat Availability

All data described in this manuscript are contained within the manuscript and supporting information. Raw data files are available upon request from the corresponding author.

