## Supplemental Figure 1 for "Utilizing a cell culture based novel cellular thermal shift assay to understand the isoform-dependent thermal stability of ApoE variants"

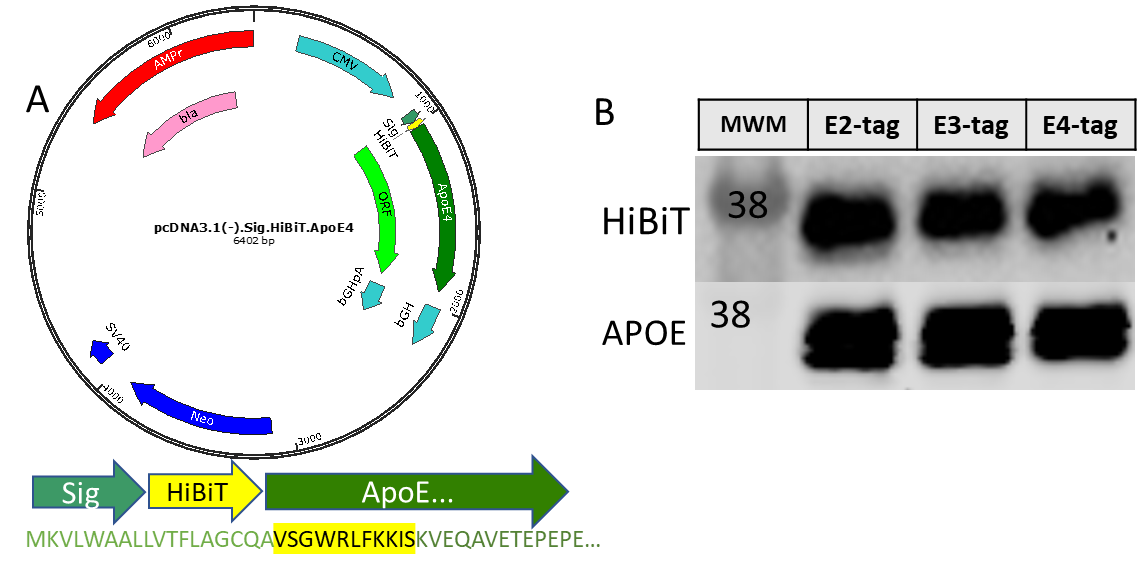

**Supplemental figure 1: Introducing the HiBiT tag onto ApoE**. A) Plasmid map for N-terminally tagged ApoE4 indicating its introduction following the signal peptide. B) HiBiT and APOE western staining for HEK cells transfected with HiBiT tagged APOE2 3 or 4.
